# Genetic variation in early-season leaf photosynthesis in sugar beet and its relationship with Cercospora leaf spot resistance

**DOI:** 10.64898/2026.04.03.716265

**Authors:** Keach Murakami, Tsubasa Narihiro, Mizuki Horikoshi, Hiroaki Matsuhira, Yosuke Kuroda

## Abstract

Improving photosynthesis is a promising approach to enhance sugar beet productivity. However, genetic variation in leaf photosynthesis and its relationship with disease resistance remain underexplored. We evaluated 98 sugar beet genotypes representing different breeding categories, including commercial F1 hybrids, seed-parent lines, and pollinator lines, in Hokkaido, northern Japan. Leaf gas exchange was measured during early growth under field conditions around the infection period of Cercospora leaf spot (CLS). To account for fluctuating irradiance during large-scale phenotyping, we applied a multilevel mixed-effects light-response model to estimate genotype-specific photosynthetic characteristics. Substantial genotypic variations in photosynthetic characteristics were detected. F1 hybrids exhibited higher photosynthetic capacity than breeding lines, whereas differences among breeding categories were unclear due to large within-category variation. Some breeding lines exhibited photosynthetic rates higher than those of hybrids, indicating exploitable genetic resources within the present genetic panel. We did not detect statistically significant trade-off between leaf photosynthesis and CLS resistance among 98 genotypes; in a subset of 19 genotypes analysed in detail, the relationship was even synergistic. Our results highlight the genetic diversity of leaf photosynthesis and its category-dependent structure, and suggest that selection for enhanced photosynthesis can proceed without substantial trade-off with CLS resistance.

**Highlight:** Leaf photosynthesis of 98 sugar beet genotypes showed significant genetic variation and dependence on breeding category. Active photosynthesis incurred minimal trade-off with Cercospora leaf spot resistance.

## 1. Introduction

Low radiation intercept has been recognized as the bottleneck of productivity in sugar beet (*Beta vulgaris* L. ssp. *vulgaris*) cultivation (Scott and Jaggard, 1993; Milford, 2006; Hoffmann and Kenter, 2018; Qi, 2022). Sugar beet intercepts sunlight and converts it into biomass via photosynthesis to accumulate sugar, providing approximately 10–20% of the world’s sugar (FAO, 2025). As long as the plants do not flower, the sugar accumulation period is indefinite and yield continues to increase (Jaggard *et al*., 2009). This simple developmental process of sugar beet highlights the importance of canopy photosynthesis. Because most commercial sugar beets are sown in spring and cover land surface slowly, a substantial fraction of solar radiation is not intercepted during spring and early summer. For example, Jaggard *et al*. (2009) pointed out that more than 30% of the solar radiation during the growth period was not intercepted by beet canopy in the United Kingdom. They also reported that warming and advanced sowing have accelerated canopy closure, which explained 68% of sugar beet yield progress during 1976–2006 (Jaggard *et al*., 2009). Earlier studies decomposed crop yield into two components: the fraction of incident radiation intercepted by the canopy (radiation intercept) and the efficiency with which intercepted radiation is converted to biomass (radiation use efficiency) (Monteith, 1977). By analyzing data from multi-genotype and multi-environment field trials, Großmann *et al*. (2026) showed that radiation interception accounted for 44% of the variation in sugar yield components, making it the most influential factor.

Another component of yield—radiation use efficiency—has been less explored than radiation intercept in sugar beet. Newly expanded leaves and canopy architecture are composed of photosynthates. Active leaf photosynthesis during the early growth phase thus enhances radiation intercept and recursively promotes canopy photosynthesis. However, there was no evidence for an improvement in radiation use efficiency in the UK sugar beets since the 1980s (Jaggard *et al*., 2009), suggesting that genetic variation for this trait remains underutilized. Free-air CO_2_ enrichment experiment demonstrated that enhanced canopy photosynthesis accelerated leaf expansion and canopy closure, resulting in 10–15 % higher white sugar yield (Manderscheid *et al*., 2010). In their multi-genotype and multi-environment data analysis, Großmann *et al*. (2026) showed that radiation use efficiency accounted for 12% of yield components. In addition, since breeding history has reduced shoot biomass relative to root biomass in sugar beet (Hoffmann and Kenter, 2018), leaf photosynthesis may become even more important in current sugar beet cultivation.

Although these studies emphasize a potential avenue to improve sugar beet productivity via promoting photosynthesis, knowledge on variations in leaf photosynthesis among genotypes is limited. Virtually all commercial sugar beet cultivars are produced via a three-way crossing scheme using seed parent lines (cytoplasmic male sterility lines and their maintainer lines) and pollinator lines (Biancardi *et al*., 2005). Because these parental lines have different prerequisite traits depending on their breeding purpose, they are expected to be functionally differentiated. Ober *et al*. (2005) measured leaf photosynthetic capacity of five hybrid cultivars and one breeding line with a genetic background of wild relatives, and found no significant differences. Bloch *et al*. (2006) evaluated leaf photosynthesis of three hybrid cultivars with different drought resistance under three watering regimes; no significant genetic variation was detected, whereas the watering regime had clear effects. In contrast, Loel *et al*. (2014) compared leaf photosynthetic rates of 17 German cultivars registered between 1964 and 2003, and found that new cultivars showed higher rates than older ones. Two recent studies compared leaf photosynthesis of several tens of Iranian sugar beet breeding lines and found significant genetic effects (Malmir *et al*., 2020; Abbasi and Bocianowski, 2021). Characterizing such diversity in leaf photosynthesis and its association with breeding category would provide a basis for exploiting the genetic resources to improve radiation use efficiency.

While improving photosynthetic performance is an attractive breeding target, it may not be without trade-offs. Active photosynthesis is connected to stomatal opening, which enables CO_2_ uptake into the intracellular space within leaf. This, however, carries the risk of disease infection because stomata are the primary entry point of phytopathogens (Wu and Liu, 2022). Cercospora leaf spot (CLS), one of the most destructive foliar diseases of sugar beet worldwide, is also caused by fungal penetration via stomata (Pool and McKay, 1916). In inoculation experiments of CLS pathogen, *Cercospora beticola* Sacc., Solel and Minz (1971) reported that resistant genotypes suppressed stomatal penetration of *C. beticola* and exhibited slower progress of CLS severity. The typical period of CLS onset is around June, which coincides with the period of active leaf expansion before canopy closure. CLS severity is influenced by the timing of epidemic onset; earlier infection leads to rapid CLS development and greater losses in final sugar yield (Yang *et al*., 2025). These raise an important question: is photosynthetic performance largely independent of CLS resistance and primarily supports yield formation, or does active photosynthesis during the early growth stage promote CLS development by increasing stomatal opening and pathogen entry? Although growth–defense trade-offs have been documented across multiple hierarchical levels in other pathosystems (He *et al*., 2022), the empirical relationship for such trade-offs in sugar beet remains open.

Sustainable management of CLS has become increasingly challenging. Current agricultural practices rely on early detection and continuous fungicide applications following CLS onset. However, fungicide-resistant strains of *C. beticola* were identified in Greece and the United States in the early 1970s (Georgopoulos and Dovas, 1973; Ruppel and Scott, 1974), and since then, resistance to a wide range of fungicide classes has been reported worldwide. Recent agricultural policies promoting reduced pesticide use further constrain chemical control options. Moreover, climate change may lead to more frequent and severe CLS outbreaks because this disease develops more rapidly under warm and wet conditions. Epidemiological modeling studies predict that under future climate scenarios, CLS onset and peak will advance in Germany, potentially causing severe damage before full canopy development is achieved (Richerzhagen *et al*., 2011; Kremer *et al*., 2016). These circumstances emphasize the central role of genetic resistance in integrated pest management strategies for CLS.

In this study, we assembled a diverse panel of sugar beet genetic resources including multigerm pollinator lines, monogerm seed parent lines (maintainers for cytoplasmic male sterility), and commercial F1 hybrids from Japanese breeding programs and international genetic resources, and addressed two questions. First, how diverse is leaf photosynthetic performance within the genetic pool, and is this variation structured by breeding category? Second, is there a quantitative trade-off between photosynthetic capacity and CLS resistance, or can the two traits vary independently? To answer these questions, we measured leaf gas exchange during the early growth stage under field conditions coinciding with natural CLS onset, and monitored CLS severity throughout the season.

## 2. Materials and Methods

### 2.1 Plant materials and climatic conditions

A three-year experiment was carried out in rainfed fields of Memuro station (42.89°N, 143.07°E) of the Hokkaido Agricultural Research Center, NARO. A panel of 98 genotypes was subjected to the analysis in the present study. This panel consisted of five hybrid F1 cultivars with known resistance and breeding lines. The five cultivars were Kawe 8K839K (KWS SAAT SE & Co. KGaA, Einbeck, Germany), Rivolta (Syngenta AG, Basel, Switzerland), Stout (SESVanderHave N.V., Tienen, Belgium), Monohikari (Hokkaido Agricultural Research Center, Japan), and Lemiel (SESVanderHave N.V.) (Table 1). Of the 93 breeding lines, 90 lines were bred at the research center, the other lines were provided from National Plant Germplasm System (Byrne *et al*., 2018; United States Department of Agriculture-Agricultural Research Service, 2025). The breeding lines consist of 16 pollinator lines and 77 seed parent lines (60 self-compatible and 17 self-incompatible lines).

**Table 1:** List of 19 genotypes subjected to core-set analysis with different Cercospora leaf spot (CLS) resistance. SI: self-incompatible, SF: self-compatible.

| Genotype | CLS resistance | Breeding category | Self-incompatibility | Origin |
| --- | --- | --- | --- | --- |
| Kawe 8K839K | Extremely resistant | F1 hybrid |  | Germany |
| Rivolta | Highly resistant | F1 hybrid |  | Switzerland |
| Stout | Resistant | F1 hybrid |  | Belgium |
| Monohikari | Medium | F1 hybrid |  | Japan |
| Lemiel | Susceptible | F1 hybrid |  | Belgium |
| NK-184mm-O | Susceptible | Seed parent | SF | Japan |
| NK-195BRmm-O | Slightly susceptible | Seed parent | SF | Japan |
| NK-197mm-O | Slightly resistant | Seed parent | SF | Japan |
| NK-207mm-O | Susceptible | Seed parent | SF | Japan |
| NK-219mm-O | Susceptible | Seed parent | SF | Japan |
| NK-235BRmm-O | Medium | Seed parent | SF | Japan |
| NK-237BRmm-O | Resistant | Seed parent | SF | Japan |
| NK-254BRmm-O | Susceptible | Seed parent | SF | Japan |
| NK-306mm-O | Resistant | Seed parent | SF | Japan |
| NK-388mm-O | Resistant | Seed parent | SF | Japan |
| NK-310mm-O | Resistant | Seed parent | SI | Japan |
| NK-303 | Resistant | Pollinator |  | Japan |
| US201 | Resistant | Pollinator |  | United States |
| FC702/2 | Resistant | Pollinator |  | United States |

Seeds were sown and raised in a greenhouse and then transplanted to open fields on 2023-05-11, 2024-05-13, and 2025-05-12. Plots were arranged in a randomized complete block design with four replicates and were maintained according to local conventional management practices. In 2023, each plot consisted of one row with four seedlings. The seedlings were spaced 22.5 cm apart within the row. The distance between adjacent plots (rows) was 120 cm. In 2024 and 2025, each plot consisted of two rows with six seedlings in total. Each row contained three seedlings spaced 22.5 cm apart. The two rows within a plot were 60 cm apart. The distance between adjacent plots ranged from 50 to 120 cm.

For sub-daily meteorological data, air temperature and solar irradiance were recorded every 10 min at the experimental station. Experimental fields were different among the three seasons but were at most 2.5 km from the observation site. Because there were some days with missing data in the observation records, we sourced daily mean air temperature and total precipitation obtained from a spatially interpolated daily meteorological dataset (Ohno *et al*., 2016; Murakami and Nagasaki, 2023) for season-scale analyses.

### 2.2 Inoculation and evaluation of CLS severity

To artificially augment natural infection, inoculation treatments were performed on 2023-07-07, 2024-07-02, 2025-07-01. Approximately 10 g of CLS pathogen inoculum was placed at the base of every plant. This inoculum consisted of soil and infected leaf debris collected during the preceding season, mixed at a dry weight ratio of 50:1. After the inoculation treatment, CLS severity was evaluated according to a local guideline (Hokkaido Agricultural Research Station, 1986). A severity of 0 indicates no spot; 1 indicates a few small spots on mature leaves; 2 indicates small spots on most mature leaves with some large lesions; 3 indicates that most mature leaves are covered with spots and partially dead; 4 indicates that several mature leaves are dead; and 4–6 indicates that most mature leaves are dead with some spots also present on newly developing leaves. Severity was rated on a 0–5 scale in increments of 0.5 as the mean of five plants per plot. The severities recorded in mid- to late-August (2023-08-15, 2024-08-29, and 2025-08-16), when differences in CLS severity among genotypes were most apparent, were averaged across the three seasons to obtain a single CLS severity score for each genotype.

### 2.3 Leaf photosynthesis measurement

During the three seasons, we evaluated leaf photosynthetic rates on five measurement days during the period of airborne pathogen exposure. Measurements were carried out around the summer solstice before the inoculation treatment except for the second measurement day in 2023 (2023-07-10) when CLS severity was zero for all genotypes. Net photosynthetic rates of leaves were measured using a portable gas-exchange measurement instrument (MIC-100-S1, Masa International Co., Ltd., Kyoto, Japan). This instrument introduces ambient air into the chamber and monitors the decrease in chamber CO_2_ concentration to calculate the photosynthetic CO_2_ uptake rate (Tanaka *et al*., 2022). The top of the chamber was transparent. Incident sunlight was used as the actinic light. The rate was calculated from the time course of the decrease in chamber CO_2_ concentration of 10–20 μmol mol^-1^ from the ambient value, which was typically completed within 5 s. Air temperature, relative humidity, and CO_2_ concentration in the chamber were not controlled.

Atmospheric CO_2_ concentrations were approximately 500 µmol mol^-1^ in the early morning and fell below 300 µmol mol^-1^ by midday owing to respiration and photosynthesis by surrounding crop fields. For each measurement, we randomly selected a mature leaf that was not shaded by neighboring individuals. In 2024 and 2025, we attached a photon sensor (DEFI2-L, JFE Advantech Co Ltd., Hyogo, Japan) adjacent to the chamber to record incident photosynthetic photon flux density (PPFD) at the measured leaf. Instantaneous PPFD was logged at five-second intervals, and moving averages were used to calculate 1-min PPFD time courses.

Measurements were repeated on two complementary sets of genotypes during the daytime under field conditions: core set and full set. The core set consisted of 19 genotypes—the five hybrid F1 cultivars and 14 breeding lines (Table 1), while the full-set scheme covered all 98 genotypes. The core genotypes were selected to cover a wide range of CLS resistance levels based on preliminary experiments. For each set, we sequentially measured a single leaf from each genotype, and this process was repeated. Measurements for the core set were conducted rapidly (approximately 10 min per sequence) to minimize fluctuations in incident radiation. The core-set scheme was repeated across measurement days, yielding a total of 69 sequences. Because a single sequence of the full-set measurement took more than one hour to complete, changes in incident radiation during the measurement period could not be ignored. We thus adopted a modeling approach considering both genotype-and environment-dependent effects as well as incidental PPFD to extract genetic variation in photosynthesis (for details, see the next section). Although we obtained 13 full-set sequences in total, only the eight sequences in 2024 and 2025, for which 1-min PPFD data were available, were used for the modeling.

In 2025, we also measured leaf stomatal conductance using a portable porometer (LI600, LI-COR Environmental Inc., Lincoln, NE, USA). The measurements were carried out immediately before or after the photosynthetic rate measurement on the same leaf at the same position. Because a positive correlation between net photosynthetic rate and stomatal conductance was confirmed (Fig. S1), photosynthetic rate serves as a good proxy for assessing stomatal openness, as reported in earlier studies (e.g., Scarth and Shaw, 1951).

### 2.4 Modeling

For analyzing full-set genotypes, leaf net photosynthetic rate was modeled as a nonlinear function of PPFD using a multilevel mixed-effects framework. The model was fitted by Bayesian regression using the brms package (ver. 2.20.4, Bürkner, 2017) with Markov chain Monte Carlo sampling in Stan (ver. 2.32.2, Stan Development Team, 2023) using R software (ver. 4.5.1, R Core Team, 2025). For the *i*-th observation of net photosynthetic rate of genotype *g* measured under environment *e*, *A_ige_* was expressed as

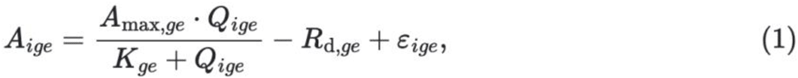

where *Q*_ige_ is the PPFD at the time of measurement, *A*_max,ge_ is the maximum photosynthetic rate, *K* is the half-saturation constant, *R*_d_ is the dark respiration rate, and *ε*_ige_ ∼ *𝒩*(0, σ^2^) represents residual error with standard deviation of σ. The three parameters were modeled as varying among genotypes and environments (i.e. sequences):

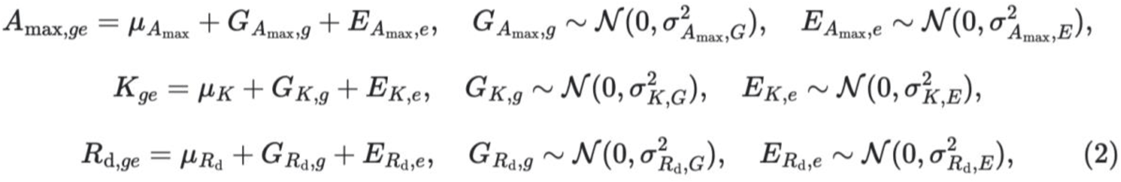

where *µ* are population-level means and *G* and *E* are genotype- and environment-specific deviations assumed to follow normal distributions. This hierarchical structure allows partial pooling across genotypes and environments, enabling robust estimation of genotype-specific photosynthetic characteristics while accounting for fluctuation in instantaneous PPFD during the sequence.

Weakly informative priors were assigned to global means of the three photosynthetic parameters to constrain estimates to biologically plausible ranges. Priors for global means of *A*_max_ and *R*_d_ (*µ*_Amax_ and *µ*_Rd_) were *𝒩*(25, 10^2^) and *𝒩*(2, 2^2^), respectively. *K* was modeled on the log scale (i.e. exp(*K*)) to ensure positive values in half-saturation. We assigned a normal prior, *𝒩*(ln(350), (ln(350)/10)^2^), corresponding to a prior median of approximately 350 µmol m⁻² s⁻¹ on the original scale. In addition, priors for genotype- and environment-specific deviations on *R*_d_ (σ*_R_*_d,*G*_ and σ*_R_*_d,*E*_) were *𝒩*(0, 0.5^2^). Default brms priors were used for the other parameters.

Convergence was assessed by confirming well-mixed trace plots of the Markov chains and small potential scale reduction factors (R-hat < 1.01). To quantify genetic differences in leaf photosynthesis under moderate light, we calculated *A*_300_ (leaf photosynthetic rates at a PPFD of 300 µmol m^-2^ s^-1^) by substituting *Q* = 300 and *E_A_*_max,e_ = *E_K_*_,e_ = *E_R_*_d,e_ = 0 in Eqns. 1 and 2. Median values of the photosynthetic characteristics calculated from the posterior distributions were used for statistical analyses.

### 2.5 Statistical analysis

The net photosynthetic rates of core-set genotypes within each sequence were standardized to allow comparison among core-set genotypes under different environmental conditions.

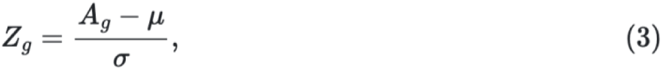

where *Ag* is the net photosynthetic rate of genotype *g*, µ and σ are mean and standard deviation of the rates measured in a sequence (i.e. 19 values). Standardized photosynthetic rates were compared among core-set genotypes using Tukey’s test. We also calculated the rank of each genotype within each sequence. Photosynthetic characteristics were compared between F1 hybrids and breeding lines, between pollinator lines and seed parent lines, and between self-compatible and self-incompatible lines using Welch’s *t*-test. Genotypes with uncertain information were excluded from these statistical tests. Pearson’s correlation coefficient was used to assess the relationship between photosynthetic characteristics and CLS severity score. Statistical significance was tested at a significance level of 5% using R software.

## 3. Results

### 3.1 CLS severity

All three seasons was characterized by record-high summer temperatures (Fig. 1). Mean air temperatures during May–July were higher than the 30-year normal values by 2.5, 2.2, and 3.6 °C in 2023, 2024, and 2025, respectively. According to local reports by Hokkaido Plant Protection Office, first occurrence of CLS in the study region were on 2023-06-27, 2024-06-26, and 2025-07-02, and the seasonal CLS incidence was classified as “moderately high” or “high” in all three years.

**Fig. 1:**
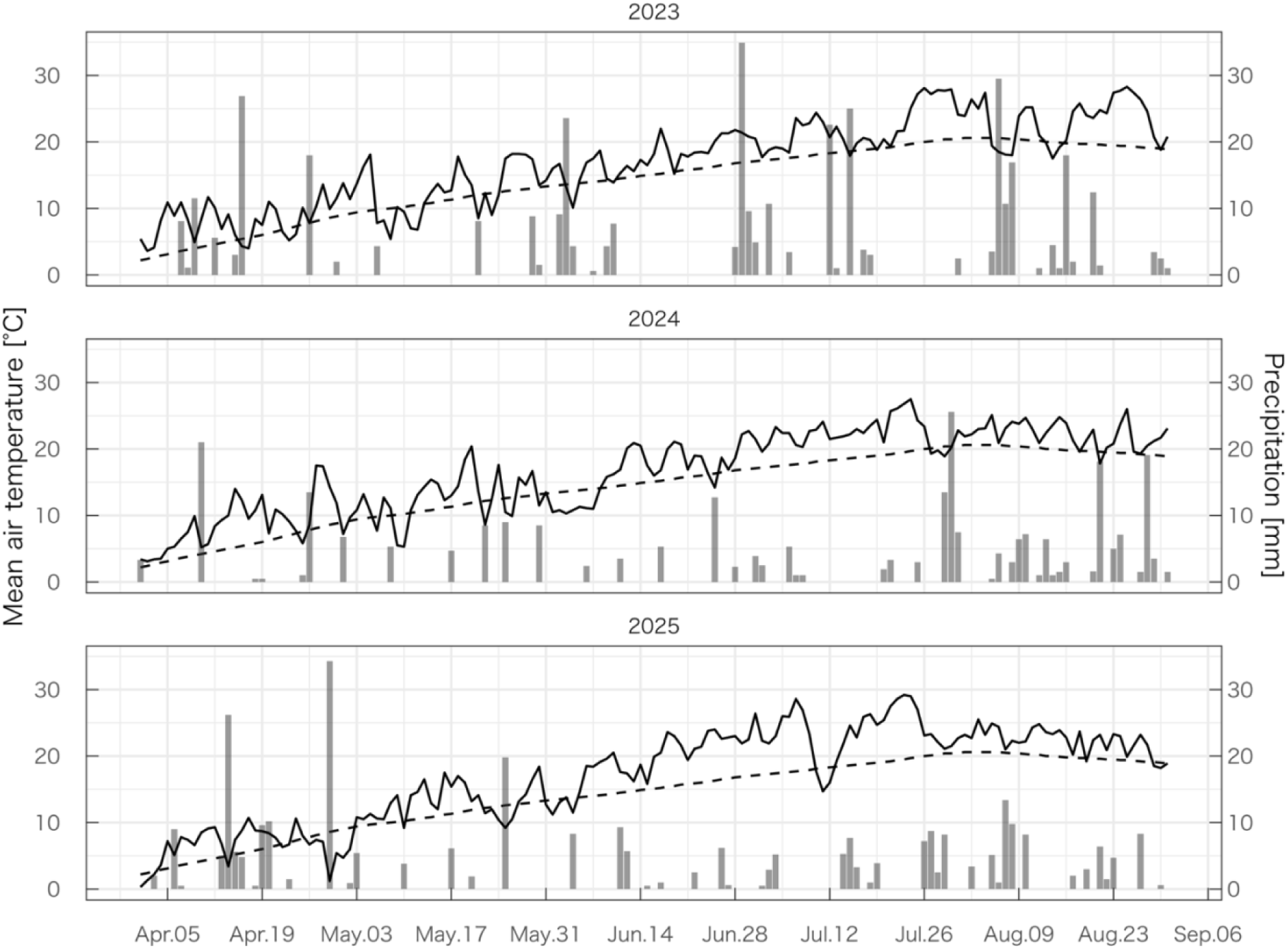
Daily average air temperatures (lines) and daily total precipitations (bars) during three years. Dashed lines indicate climatological normal air temperatures (1991–2020).

The progress of CLS severity differed among the three seasons in the experimental fields, presumably reflecting differences in weather conditions (Fig. 2). In 2023, wet conditions from late June through July caused a steep increase in severity after CLS onset. In 2024, relatively dry conditions in June and July slightly delayed CLS onset, but severity increased steadily thereafter. In 2025, warm temperatures in June combined with intermittent rainfall resulted in high severity already by late July. CLS severity of genotypes showed generally consistent trends among three seasons (Fig. 2 inset). The severity of five F1 hybrids with known resistance levels followed approximately the order previously reported: Kawe 8K839K showed the highest resistance, followed by Rivolta, with Lemiel showing the weakest resistance.

**Fig. 2:**
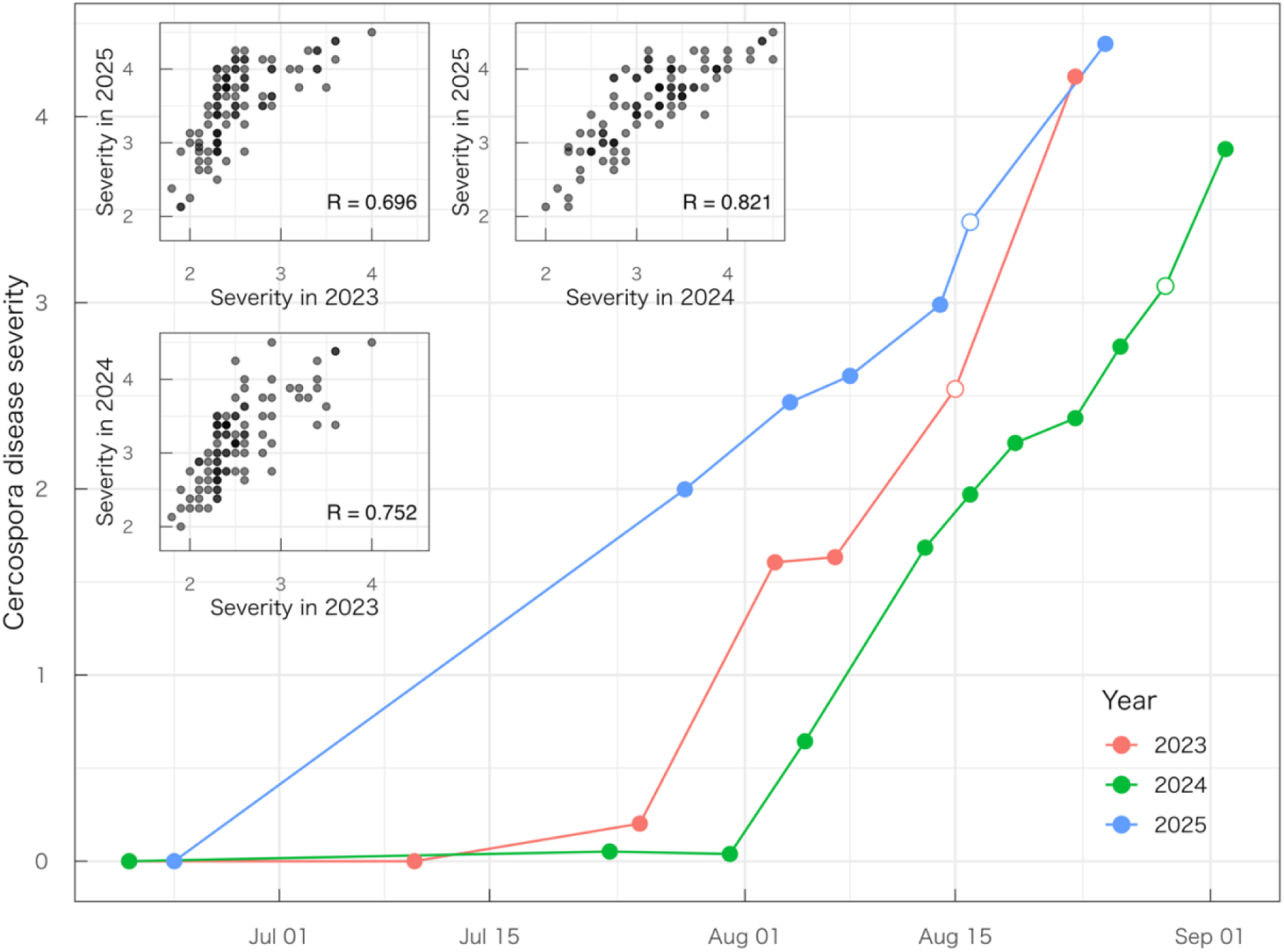
Progress of Cercospora disease severity during three years. Mean severity values of 98 genotypes are shown. Open symbols indicate data used for calculating 3-year average severity score for each genotype. Inset shows pairwise correlations of the severity among 98 genotypes between two seasons.

### 3.2 Genetic variation of leaf photosynthesis

Leaf net photosynthetic rate followed the course of solar irradiance, except on 2025-06-24, when leaf photosynthesis appeared to be suppressed by heat and/or drought stress (Fig. 3). The rate showed substantial difference within each sequence of measurements, suggesting genetic differences in photosynthetic characteristics.

**Fig. 3:**
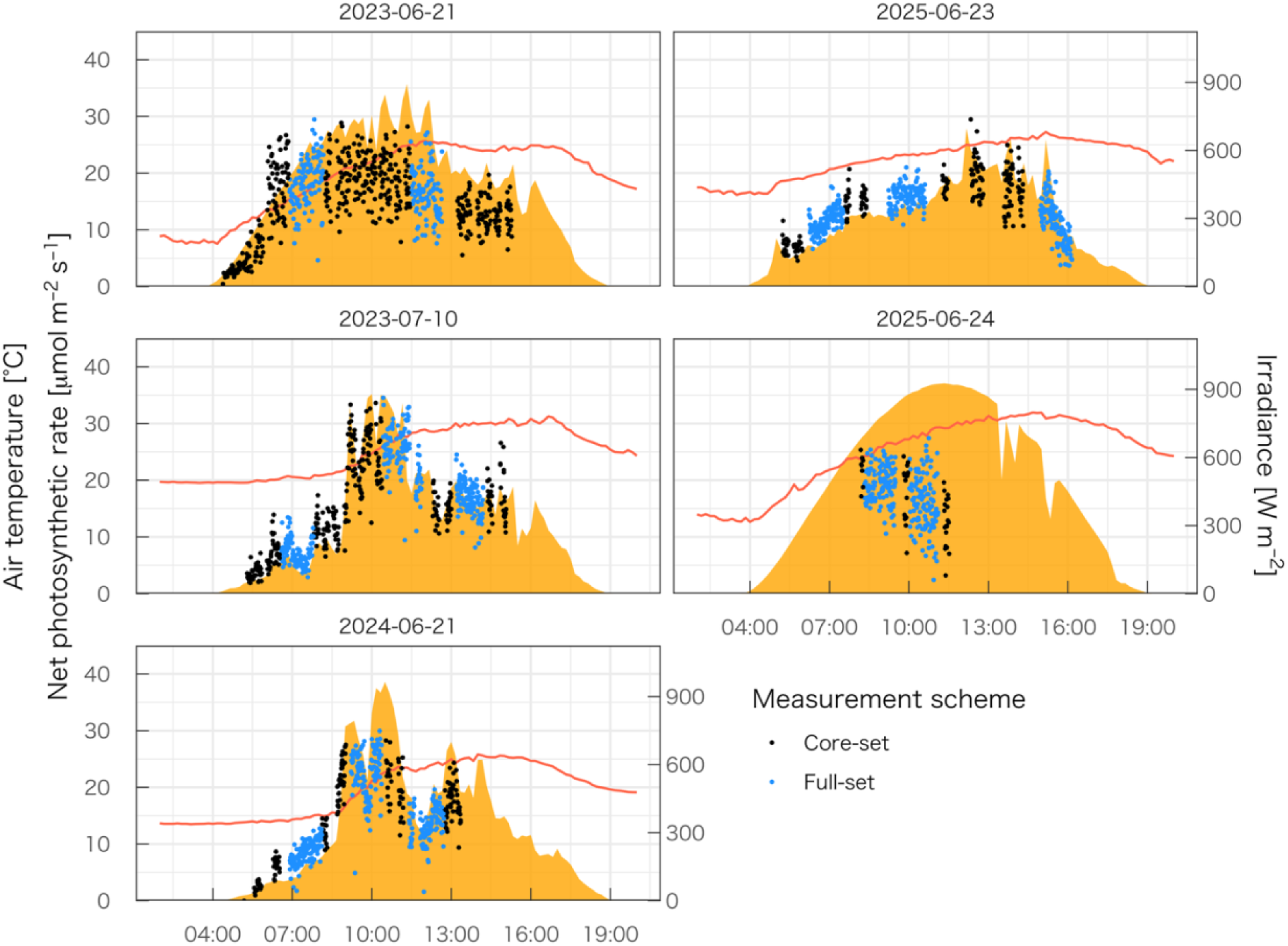
Diurnal time courses of 10-minute average air temperature (lines), solar irradiance (areas), and net photosynthetic rate (points) of sugar beet leaves.

Daily averaged leaf photosynthetic rates of the core-set genotypes were positively correlated among five measurement days (Fig. S2A). This indicates that our scheme successfully and consistently captured genetic variation.

Standardized photosynthetic rates differed among the core-set genotypes (Fig. 4A). The standardized rates were higher in the five F1 hybrids than in the breeding lines. Among the hybrids, resistant cultivars (Kawe 8K839K and Rivolta) showed higher standardized rates, whereas Stout, Monohikari, and Lemiel showed rates comparable to those of the breeding lines. The standardized rates were notably low in some breeding lines (e.g. NK-388mm-O, NK-235BRmm-O, and NK-207mm-O). The resistant cultivars, Kawe 8K839K and Rivolta, consistently showed high ranks in the net photosynthetic rate among 19 genotypes irrespective of the sequence (Fig. 4B). The ranking of breeding lines imported from the United States (FC702/2 and US201) varied widely depending on the sequence (Fig. 4B).

**Fig. 4:**
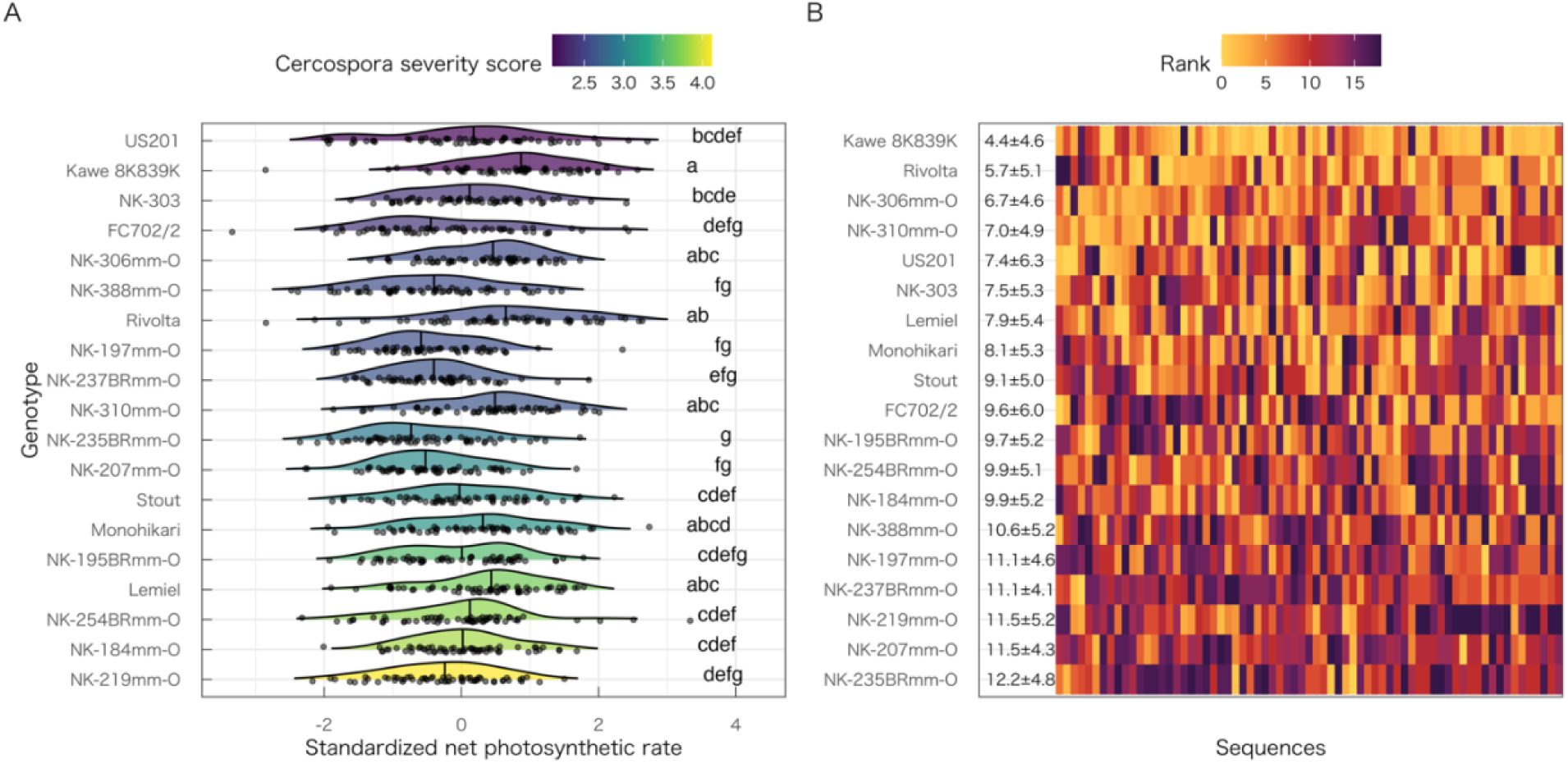
(A) Genetic variation of the distributions of standardized net photosynthetic rates. Small dots and vertical bars indicate observed values and their medians (N = 69). The rates of genotypes with different lower case letters were statistically different (P < 0.05). (B) Genetic variation of the ranks in the net photosynthetic rate among 19 genotypes under 69 different sequences. Means ± standard deviations of the rank are shown. Genotypes are sorted according to the disease severity and the mean rank in panel A and B, respectively.

In contrast to the core genotypes, there were weak or no correlation in daily averaged leaf photosynthetic rates of full-set genotypes among the five measurements days (Fig. S2B), perhaps due to fluctuating irradiance during a sequence. This was because the full-set measurements required substantially longer time per sequence (> 1 h vs. ∼ 10 min for the core-set sequence). To account for this fluctuating irradiance, we adopted a multilevel mixed-effects modeling framework. The model estimated genotype- and environment-specific photosynthetic light-response curves across the range of observed photon flux densities (Fig. S3). For each environment (i.e., each of eight full-set sequences), our model captured light-saturating response of net photosynthetic rate while allowing for systematic variation among genotypes. The estimated net photosynthetic rates closely matched the observed values across environments (Fig. 5), indicating that the model properly reproduced measured photosynthetic rates under field conditions.

**Fig. 5:**
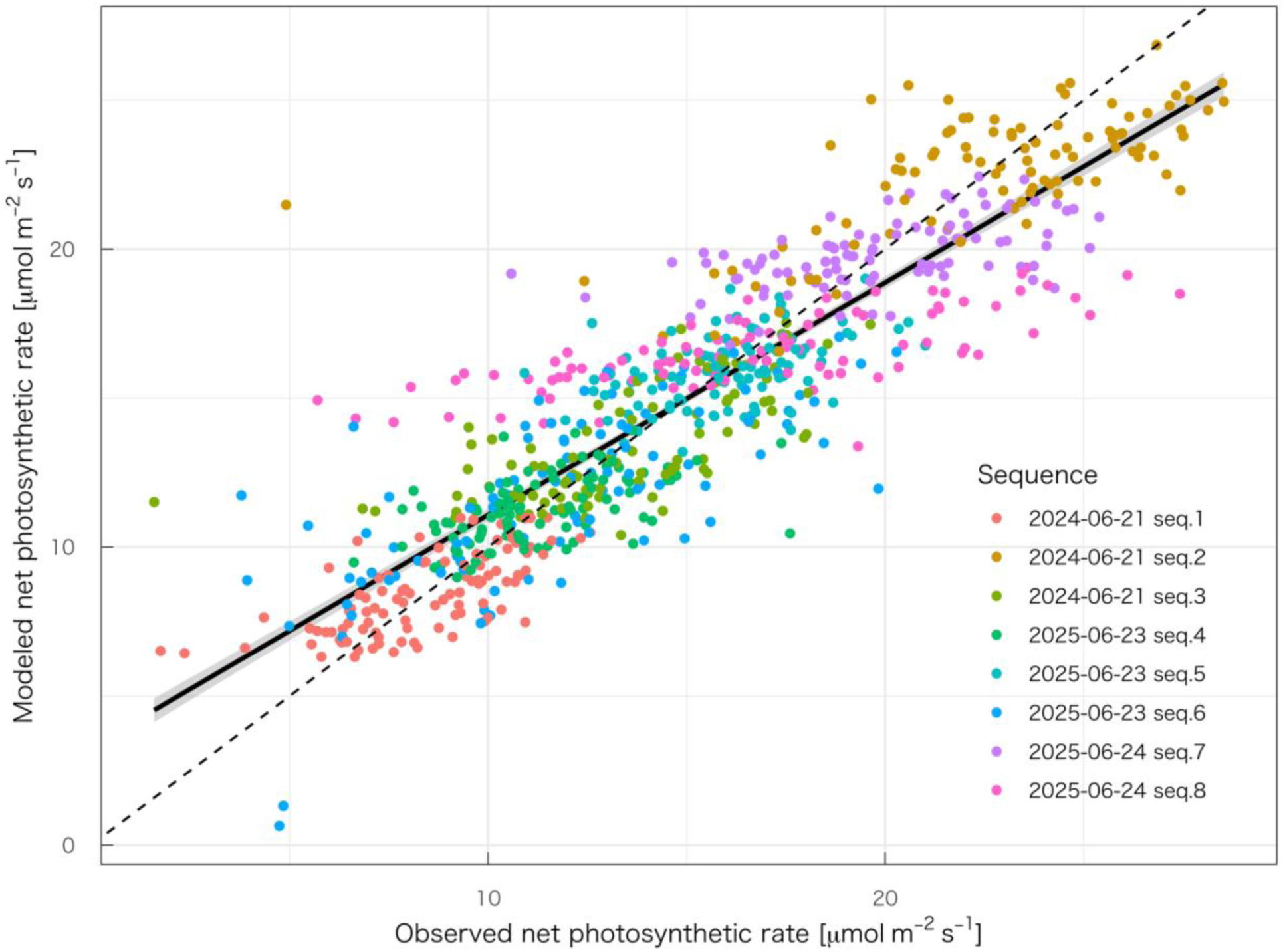
Relationship between observed and model-estimated leaf photosynthetic rates of sugar beet.

### 3.3 Leaf photosynthesis and breeding category

The maximum photosynthetic rate (*A*_max_) estimated by the model was significantly higher in F1 hybrids than in breeding lines (Fig. 6). *A*_max_ tended to be higher in pollinator lines than in seed parent lines, although this difference was not statistically significant. There was no statistical difference in *A*_max_ between self-compatible and self-incompatible seed parent lines. Note that some self-compatible lines showed substantially low *A*_max_. Net photosynthetic rate under moderate light (*A*_300_) and dark respiration rates (*R*_d_) showed similar patterns to *A*max while the half-saturation constant (*K*) showed no significant difference among breeding categories (Fig. S4).

**Fig. 6:**
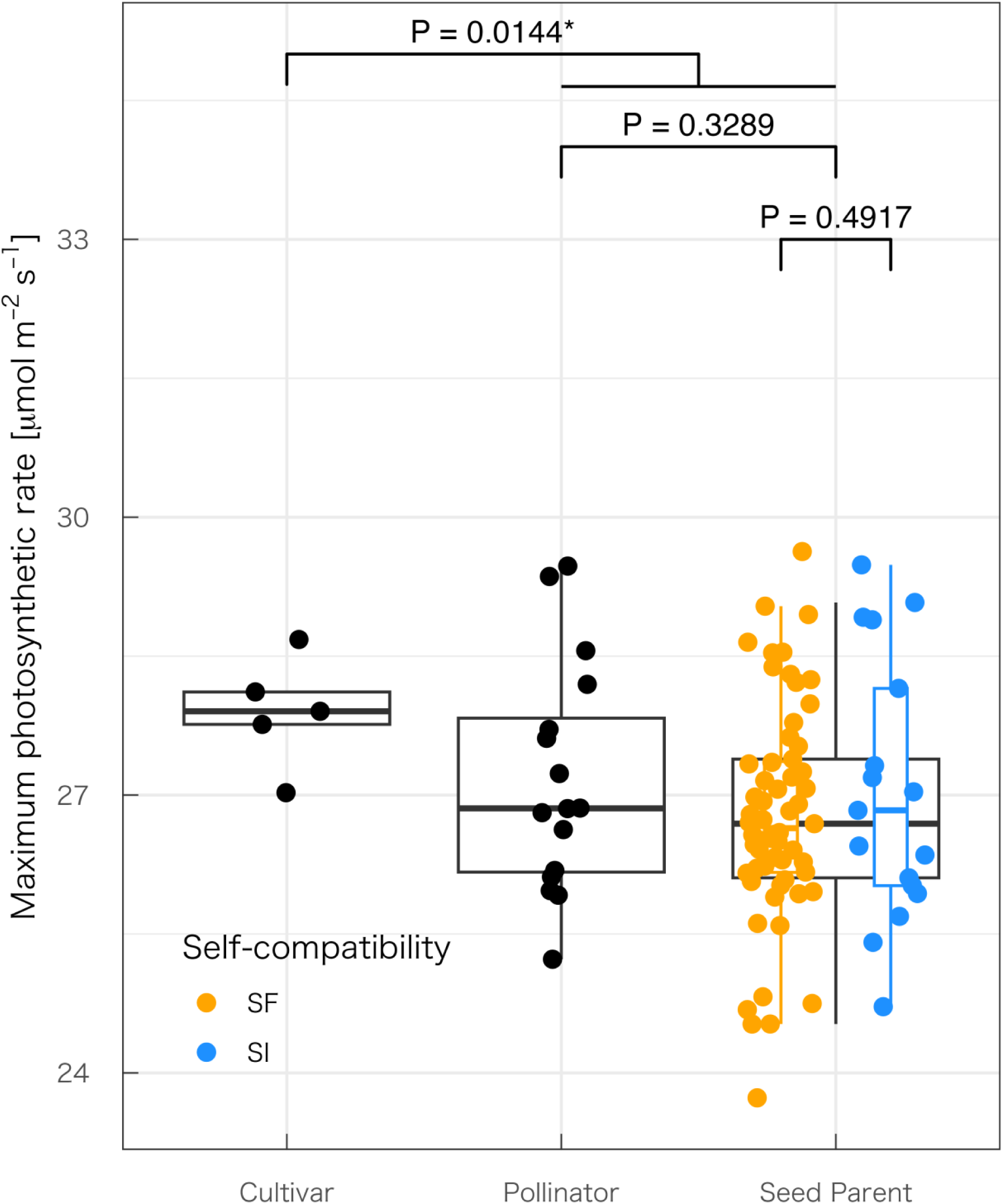
Maximum photosynthetic rate of sugar beet leaves of F1 hybrids, pollinator lines, and seed parent lines. For seed parent lines, boxplots of self-compatible (SF) and self-incompatible (SI) lines are separately shown with their aggregated result (wider black one).

### 3.4 Leaf photosynthesis and CLS severity score

For core-set genotypes (Fig. 7), correlation between the standardized photosynthetic rate and CLS severity score was negative but not significant (R = −0.18, P = 0.462). For full-set genotypes (Fig. 8), there were no statistical correlation between CLS severity score with net photosynthetic rate under moderate light (*A*_300_; R = 0.07, P = 0.513) and that under saturating light (*A*_max_; R = 0.09, P = 0.398). No significant correlation was detected for the half-saturation constant *K* and the dark respiration rate *R*_d_ (Fig. S5).

**Fig. 7:**
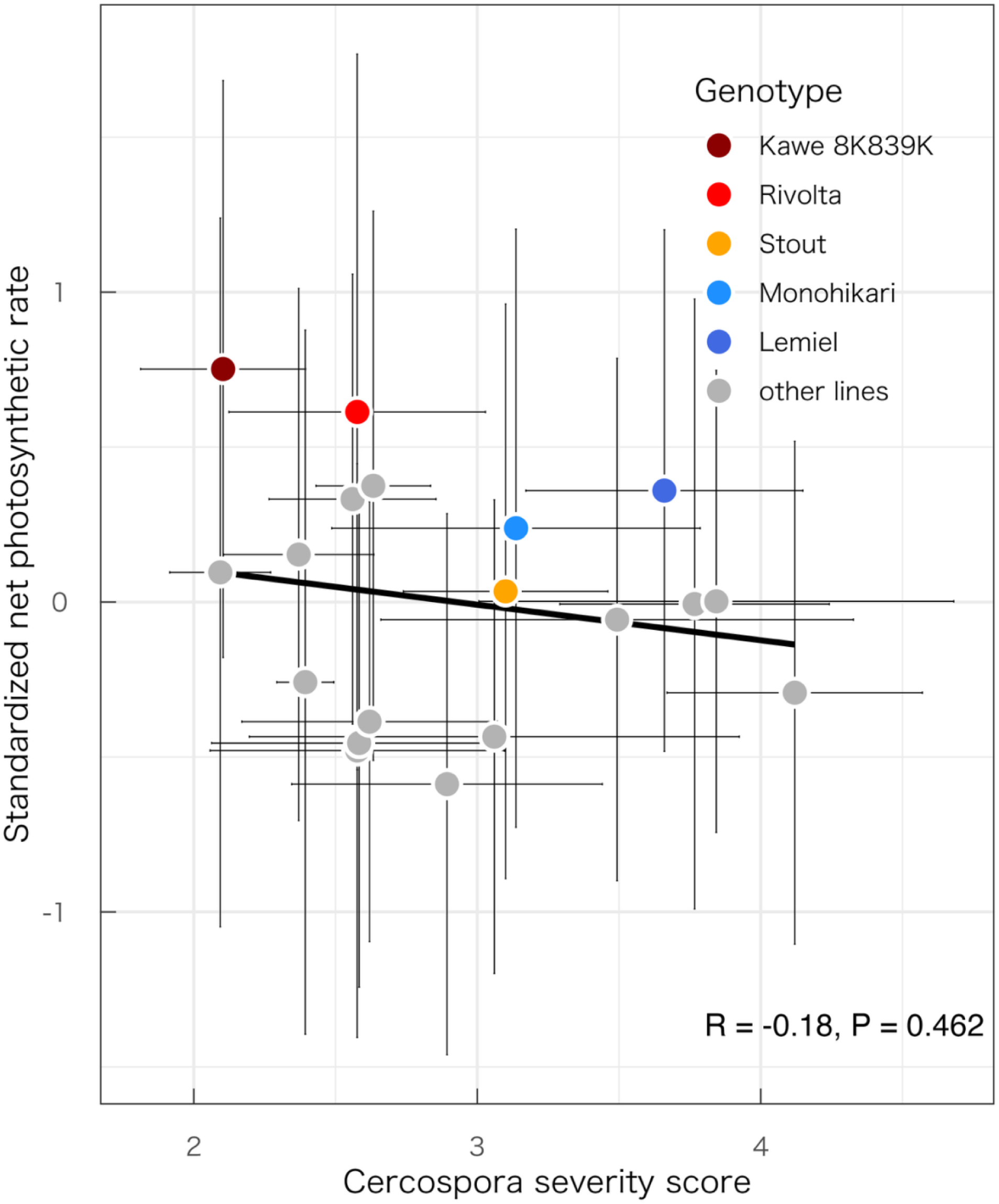
Relationship between standardized net photosynthetic rates and Cercospora severity scores of 19 sugar beet genotypes. For the photosynthetic rates and severity, means ± standard deviations of 69 sequences and 3 years are shown, respectively.

**Fig. 8:**
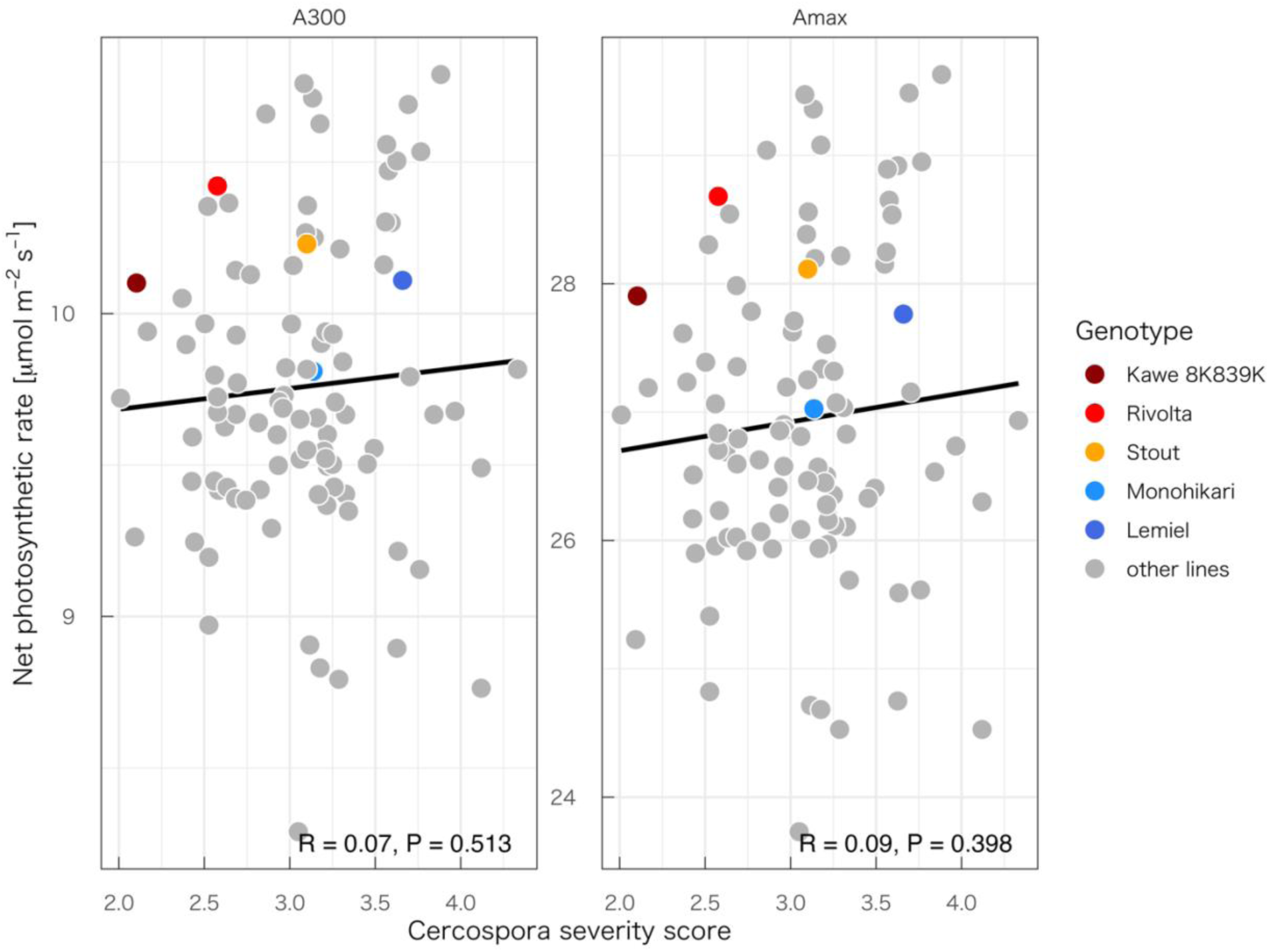
Relationship between photosynthetic rates under moderate light (*A*_300_) or saturating light (*A*_max_) and Cercospora severity scores of 98 sugar beet genotypes. For the photosynthetic rates and severity, medians of posteriors and 3 years are shown, respectively.

## 4. Discussion

### 4.1 Genetic variation for improving photosynthesis

Improving photosynthesis by exploiting genetic resources is a promising approach to enhance sugar beet productivity. Recent genome-wide association analyses on sugar beet have explored genomic effects on various agronomic traits (Würschum *et al*., 2011; Li *et al*., 2023; Liu *et al*., 2023; Wang *et al*., 2023; Wang *et al*., 2024). However, photosynthetic characteristics are not included despite their importance on sugar productivity. This gap may be related to low throughput of photosynthesis measurements. In addition, the light response of photosynthesis is highly dynamic over time and measuring environments, easily masking genetic effects on photosynthesis (Fig. S2B).

Here, we addressed these challenges by applying two schemes. In the core scheme, intensive gas exchange measurements using a high-throughput device (Tanaka *et al*., 2022) successfully revealed genetic variation in leaf photosynthesis under field conditions. In addition, rank analysis within the core-set genotypes demonstrated the sensitivity of leaf photosynthesis to the measurement environment differed among genotypes. The US-origin lines whose ranking varied depending on the measurement sequence may be sensitive to environmental factors such as temperature and drought. In the full-set scheme, we applied multilevel modeling to separate effects of genotype, environment, and instantaneous irradiance. This method quantified genetic variation of photosynthesis across a large population. The approach combining intensive field phenotyping and statistical modeling has been applied in breeding research (e.g. Keller *et al*., 2024; Gao *et al*., 2025), and the present results further emphasize its utility.

As a benefit of comparing multiple genotypes, our analysis revealed the structure of leaf photosynthesis depending on breeding categories (Fig. 6). Commercial F1 hybrids showed higher photosynthetic capacity than breeding lines in the full-set analysis, which was observed but less clearly in the core-set analysis. This was presumably due to heterosis in photosynthesis, reported in some species (Fujimoto *et al*., 2012; Zhu *et al*., 2020) as well as sugar beet (Usui *et al*., 2022). Greater biomass accumulation in commercial F1 hybrids than in breeding lines, which is usually observed in sugar beet, can be explained by the improvement in leaf photosynthesis. Because F1 cultivars have been subjected to strong selection pressure for biomass and sugar productivity, photosynthetic capacity may have been indirectly screened during breeding trials through its association with biomass accumulation. Note that biomass accumulation at the crop level is determined by both leaf-level photosynthetic characteristics and canopy architecture (i.e. LAI and vertical light profile; Monsi & Saeki, 1953). The present study focuses on genetic variation in the former component, and future work should integrate LAI to evaluate their combined effects on biomass and final sugar yield. The difference in photosynthetic capacity was less obvious between pollinator lines and seed parent lines or between self-compatible and self-incompatible lines. This was due to large variation in photosynthetic capacity of breeding lines (Fig. S6), which reduced statistical power to detect significant effects. There were some breeding lines that outperformed hybrids in photosynthesis, implying that beneficial genetic resources for photosynthesis are present within our population. Among different breeding categories, self-compatible seed parent lines showed largest variation in photosynthetic capacity. Because self-compatible seed parent lines have been maintained by selfing, leaf photosynthesis of some of these lines may be suppressed due to inbreeding depression. Our breeding lines used in this study originate from a limited pool of genetic diversity (approximately 10 open-pollinated varieties) (Taguchi et al., 2019). These differences in prerequisite traits and maintenance methods among the breeding categories might shape diversity of leaf photosynthesis in the present study. The present analysis, which links breeding information with photosynthetic traits, enables inference that the potential for enhancing photosynthesis may be greater on the seed parent side, thereby providing guidance for the direction of breeding programs.

### 4.2 Active photosynthesis incurs minimal resistance cost to CLS

The second research question of this study was whether enhanced leaf photosynthesis—and the concomitant stomatal opening—promotes CLS development by facilitating pathogen entry through stomata. The positive correlation between net photosynthetic rate and stomatal conductance (Fig. S1) indicates that genotypes with higher photosynthetic rates maintain stomata open during the period of airborne pathogen exposure. However, in full- and core-set analyses, we did not detect statistically significant correlation between photosynthetic capacity and CLS severity. We thus did not observe a growth–defense trade-off in the present system. Molecular studies in model plants have revealed that growth–defense trade-offs is mediated by hormone signaling crosstalk and transcription factor balancing, implying an inevitable relationship under precise regulatory control (Ning *et al*., 2017). In crop systems, the trade-off remains unclear regarding both the existence and the magnitude (Brown, 2002). Nevertheless, the absence of correlation between leaf photosynthesis and CLS severity implies that stomatal openness is not a dominant factor of CLS resistance. Downstream molecular resistance mechanisms play a major role. The search for molecular resistance loci—such as the recently identified BvCR4 resistance gene (Törjék *et al*., 2023)—remains a critical priority.

While a trade-off between photosynthetic activity and CLS resistance cannot be entirely dismissed, the present results indicate that breeding can achieve simultaneous improvement in both traits. Moreover, genotypes with higher resistance tended to exhibit higher photosynthetic capacity in the core-set analysis. Because the net photosynthetic rate of core-set genotypes estimated by applying the model was consistent trends in response to CLS severity score (Fig. S7), this synergistic result was not attributable to the method. In sugar beet, the yield penalty associated with CLS resistance has diminished progressively over the breeding history. Resistant cultivars showed approximately 18% lower sugar yield than moderately resistant ones in Midwestern US trials in the early 1990s (Miller *et al*., 1994). This yield penalty was 5–7% in German trials with cultivars released in the mid-2000s (Gummert *et al*., 2015). Among German cultivars released in the 2010s, no detectable yield penalty was observed (Vogel *et al*., 2018). In breeding programs of Japanese sugar beet, we have recently developed highly CLS resistant candidates with high yield potential (unpublished data). Our results are consistent with this historical trajectory, indicating that the decoupling of productivity and CLS resistance has progressed to the point where vigorous early-season photosynthesis incurs a minimal resistance penalty.

The sugar beet–*C. beticola* pathosystem has a long history in stomatal immunity: it was among the first systems in which fungal penetration through stomata was documented over 140 years ago (Wu and Liu, 2022). Pool and McKay (1916) confirmed through microscopic examination of fixed specimens that penetration occurs exclusively through open stomata. They also found that lesions are absent on leaves whose stomata have not yet reached a certain developmental size (Pool and McKay, 1916). Subsequent work showed that germ tubes elongate randomly regardless of stomatal aperture, but that appressorium and infection peg formation require detection of open stomata and are therefore suppressed under conditions promoting stomatal closure, such as high humidity (Rathaiah, 1977) and darkness (Narisawa, 1972). Comparisons of stomatal penetration between resistant and susceptible genotypes have yielded mixed results: Solel and Minz (1971) reported lower penetration rates in resistant genotypes, whereas Narisawa (1972) found no such relationship. These earlier studies, conducted from a pathological perspective, did not quantify stomatal aperture. The genetic variation in stomatal gas exchange documented in the present study may partly account for such discrepancies. However, because we used photosynthetic rate as a proxy for stomatal openness, more direct stomatal phenotyping—for example, thermal imaging of transpiration cooling—combined with resistance evaluation would be needed to resolve this question.

### 4.3 Conclusions

By combining high-throughput field gas exchange measurements with multilevel modeling, we evaluated genetic variation in leaf photosynthesis among 98 sugar beet genotypes in different breeding categories, including commercial F1 hybrids, pollinator lines, and self-compatible and self-incompatible seed parent lines. Our results demonstrate that (1) wide genotypic variation in leaf photosynthetic characteristics existed within the present genetic panel; (2) commercial F1 hybrids exhibited higher photosynthetic capacity than breeding lines, likely reflecting heterosis and indirect selection for productivity; and (3) among breeding categories, self-compatible seed parent lines showed the largest variation in photosynthetic capacity, suggesting that this breeding category may offer the greatest scope for photosynthetic improvement. Furthermore, statistically significant relationship was not found between leaf photosynthesis and CLS resistance in the full- and core-set analyses. This indicates that active early-season photosynthesis—and the associated stomatal opening—incurs minimal resistance costs, and that simultaneous improvement of both traits is feasible through breeding.

## Supplementary data

Fig. S1: Relationship between net photosynthetic rate and stomatal conductance. Fig. S2: Correlation matrices of daily mean net photosynthetic rates.

Fig. S3: Photosynthetic light response curves estimated by the multilevel model. Fig. S4: Effects of breeding category on photosynthetic characteristics.

Fig. S5: Relationship between photosynthetic characteristics and Cercospora severity score.

Fig. S6: Estimated maximum photosynthetic rate of parental lines.

Fig. S7: Relationship between net photosynthetic rate under moderate light estimated by the multilevel model and Cercospora severity score in core-set genotypes.

## Author contributions

KM: conceptualization; KM and TN: methodology; KM and TN: formal analysis; KM, TN, and MH: investigation; TN, HM, and YH: resources; KM and TN: data curation; KM: writing - original draft; TN, MH, HM, and YK: writing - review & ediging; KM: visualization; HM and YK: supervision; KM, TN, MH, HM, and YK: funding acquisition

## Data availability

Daily meteorological data are publicly available at the Agro-Meteorological Grid Square Data system, NARO (https://amu.rd.naro.go.jp/wiki_open/doku.php). Several plant materials used this study were provided from USDA by permission. Data will be shared on request to the corresponding author with permission of USDA.

